# Histone lactylation: A new epigenetic axis for host-parasite signalling in malaria?

**DOI:** 10.1101/2022.09.22.509018

**Authors:** Catherine J. Merrick

## Abstract

Epigenetic marks such as histone acetylation and methylation play important roles in the biology and virulence of malaria parasites. Here I report that histone lactylation is also found in these parasites, and speculate on its potential functions. Lactylation is a new epigenetic modification, discovered only in 2019 in human cells. This nascent field has since focussed largely on human biology, but lactyl epigenetic marks could be particularly important in malaria parasites, which are exposed to high and fluctuating lactate levels in their host environment. This is because parasites in the bloodstream respire by glycolysis, producing lactate, and hyperlactataemia is characteristic of severe malarial disease. Therefore, blood lactate could be a signal for the status of the infected host, which could be directly translated to virulence responses via histone lactylation and modulation of parasite gene expression. Responses could include the rate of conversion into sexual transmission stages, the expression of cytoadherence genes – which enhance immune evasion by the parasite but can exacerbate pathology in the host – and the modulation of parasite stress-resistance. Lactylation may soon join acetylation and methylation as a key tool in the epigenetic arsenal of *Plasmodium*.

## MAIN TEXT

Epigenetic phenomena control many aspects of gene expression. Epigenetic changes can be fast, flexible and reversible, making them ideal for responding to changing environments. In the malaria parasite *Plasmodium*, epigenetic marks control virulence pathways including antigenic switching and alternate invasion pathways [1]. Classical histone marks like acetylation and methylation, plus their protein ‘writers’ and ‘readers’, have all been identified [2]. Thus, although *Plasmodium* genomes are unusual in many ways, they apparently make conventional use of epigenetics to control gene expression, particularly where rapid responses to varying host conditions can be beneficial. In fact, epigenetics may be particularly prominent in malaria parasites, which encode an unusual paucity of specific transcription factors.

A recent publication described the lactylation of histone lysine residues as a novel epigenetic feature in mammalian cells [3]. 28 lactylated lysines (KLa) in 4 core histones were detected (Fig 1A). The modification was dependent on lactate levels, altered either by adding extracellular lactate to cultured cells, or by stimulating intracellular glycolysis. This exciting report immediately prompted the question of whether lactylated histones might exist in *Plasmodium*, which often causes hyperlactataemia in its mammalian hosts, thus exposing itself to high and fluctuating lactate levels.

**Figure 1:**
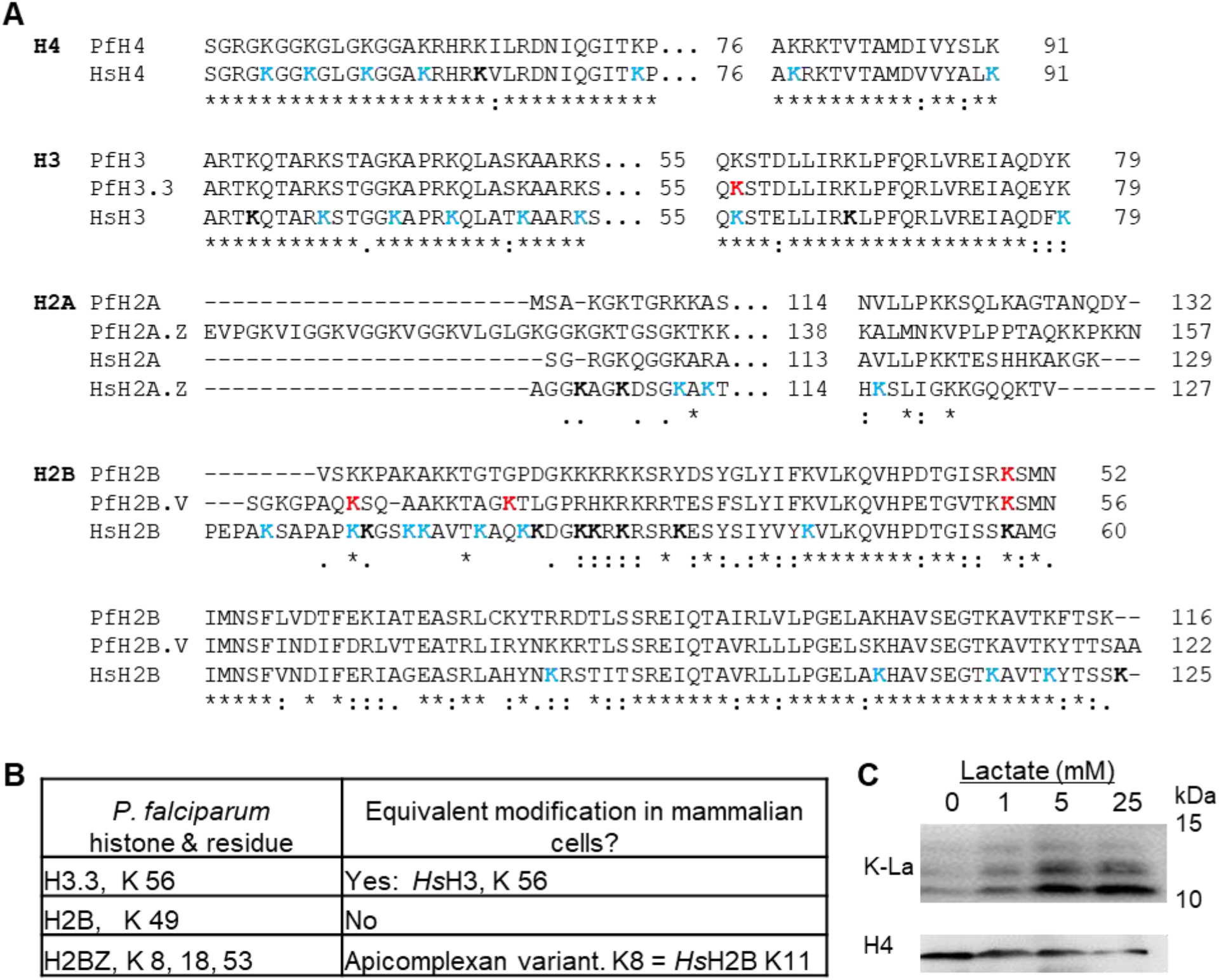
Evidence for histone lactylation in *P. falciparum* parasites A) Alignments of histones from *P. falciparum* and *H. sapiens*. KLa sites identified in human cells [3] are highlighted in blue, and those in *P. falciparum* in red. B) Table of KLa sites identified in *P. falciparum* histones in one or more of six independent mass spectrometry datasets, on protein extracted from normal *in vitro* cultures (no added lactate). C) Western blot with pan-KLa antibody on *P. falciparum* histones from trophozoite-stage parasites after 16h of exposure to increasing levels of added lactate. Total histone H4 is shown as a control.

I therefore examined mass spectrometry datasets from laboratory-cultured *P. falciparum* parasites. (Mass spectrometry was previously conducted on *P. falciparum* histones to detect acetylated residues [4], but lactylated residues were not reported – nor, presumably, sought in this 2006 analysis). I detected a characteristic shift of 72.021 Da on histone-derived peptides: 5 were detected on 3 parasite histones (Fig 1B). Therefore, *P. falciparum* does indeed have histone lactylation. These parasites had been cultured normally without added lactate, so only a subset of the most abundant modifications would probably be detected. Western blots with a pan-KLa antibody that was previously used in human cells [3] further showed that parasite histone lactylation was inducible (Fig 1C).

### Lactate and severe malaria

Hyperlactataemia is a cardinal feature of *P. falciparum* malaria [5] and a strong predictor of severe disease [6]. It correlates with parasite load, which likewise predicts severe disease. Its aetiology (recently reviewed in [7]) includes glycolysis by parasites and also by human tissues when normal oxygenation is impeded by parasitized cells sequestered in capillaries. The result is potentially fatal metabolic acidosis and respiratory distress. The clinical threshold for hyperlactataemia, 5mM, is readily reached in malaria patients. In a study conducted in 2009 [8], I reported that 29 of a cohort of 109 Gambian patients exceeded this level, reaching a maximum of 15.3mM blood lactate (Fig 2). *In vitro*, lactate at and above the 5mM clinical threshold clearly induces lactylation of parasite histones (Fig 1C).

**Figure 2:**
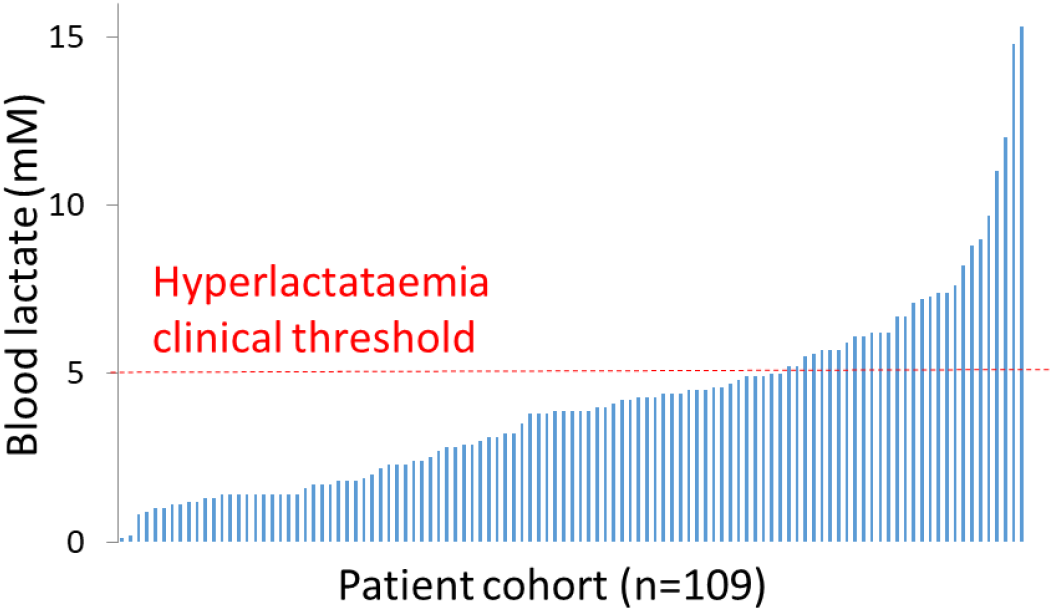
Blood lactate in malaria patients frequently exceeds a 5mM threshold Blood lactate levels (rank-ordered) measured in 109 Gambian malaria patients. Data re-plotted from [8].

### Proposed effect of histone lactylation in *Plasmodium*

Since hyperlactataemia correlates with parasite load and also with severe disease, it could simultaneously provide a crude ‘quorum sensor’ and a measure of host stress. Malaria parasites might have usefully evolved to respond to these signals by altering their growth and virulence.

In the above-referenced Gambian patient study, we found that hyperlactataemia correlated, in the causative parasites, with high expression of virulence genes in the *‘var’* family and also of their regulators – histone deacetylase enzymes called sirtuins. Therefore, we originally proposed that the parasites might sense hyperlactatemia and then activate histone deacetylation to modulate epigenetically-controlled *var* genes [8]. However, new evidence of histone lactylation now supports a simpler theory: hyperlactataemia could directly affect the epigenetic control of *Plasmodium* virulence genes, by causing histone lactylation within those genes (rather than acting ‘indirectly’ through sirtuin induction and altered histone acetylation). Lactylated histones in human leukocytes do indeed stimulate gene transcription, similar to the well-characterised effect of acetylated histones [3]. Since the basic tenets of epigenetics are largely conserved even in this early-diverging eukaryote, it is probable that lactylation stimulates transcription in *Plasmodium* as well.

Interestingly, 4 out of 5 lactylated residues found in *Plasmodium* thus far (Fig 1) were in variant histones – H3.3 and H2BZ – not in standard core histones H2A, H2B, H3 and H4. If histone lactylation is particularly abundant in variant histones, this would be significant because they are known to be associated with active genes, particularly active virulence genes [9, 10]. For example, both H2BZ and H3.3 are found in the promoters of active but not inactive *var* genes, and H2BZ-containing nucleosomes tend to be acetylated [9, 10]; Figure 1 now suggests that they are also lactylated.

### Biological implications of histone lactylation in *Plasmodium*

In some circumstances it might be beneficial for parasites to upregulate virulence genes in stressed hosts experiencing hyperlactataemia. For example:

A) Elevated expression of *var* genes can help parasites to cytoadhere to epithelial cells, sequestering them from the blood flow to avoid splenic clearance and thus replicate more efficiently [11].

B) Increased conversion to sexual gametocytes (gametocytogenesis) is beneficial in promoting mosquito-borne transmission from stressed hosts. Gametocytogenesis is another virulence phenotype that was linked to lactate in a recent publication: when parasites were cultured in clinically-achievable levels of added lactate, they converted at a higher rate [12]. This conversion requires a suite of gametocyte-specific, epigenetically-controlled genes, including the *Plasmodium*-specific transcription factor AP2-G [13], a clear candidate for regulation via histone lactylation.

C) It could be beneficial for parasites in hyperlactataemic hosts to upregulate stress-resistance genes, since such hosts are prone to severe disease with associated inflammation and fever exerting high levels of oxidative stress and thermal stress on the causative parasites. Indeed, we have generated preliminary evidence [14] suggesting that moderate lactate exposure can improve stress-resistance in cultured parasites. Genes that might be induced here are unknown but antioxidant and chaperone pathways are likely candidates. Again, such pathways could, *in vivo*, improve parasite survival in a stressed hyperparasitaemic host. Overall, lactyl epigenetic marks clearly have potential to affect parasite virulence in multiple ways.

### What is the molecular mechanism of histone lactylation?

Epigenetic marks are usually dynamic, added and removed by opposing enzymes, e.g. histone acetyltransferases and deacetylases (HATs/HDACs). Histone lactylation enzyme(s) have yet to be fully identified in mammalian cells but there is evidence for the HAT p300 [3, 15]. This lacks a direct homologue in *Plasmodium* and other GNAT-family HATs may perform its functions [16]. *Plasmodium* has a relatively small HAT/HDAC repertoire [16, 17], only some of which are experimentally confirmed. There is 1 MYST-family and up to 10 putative GNAT-family acetyltransferases, but only 2 have been fully characterised as HATs and most are probably not histone-directed. There are five HDACs, most of them confirmed as genuine HDACs, including the sirtuins mentioned above [8]. Table 1 highlights the enzymes experimentally reported as essential or inessential in erythrocytic *Plasmodium* parasites. Some HATs/HDACs could plausibly ‘moonlight’ upon lactyl as well as acetyl modifications, although it is theoretically possible that *Plasmodium* has unique lactylation enzymes as well.

**Table 1:**
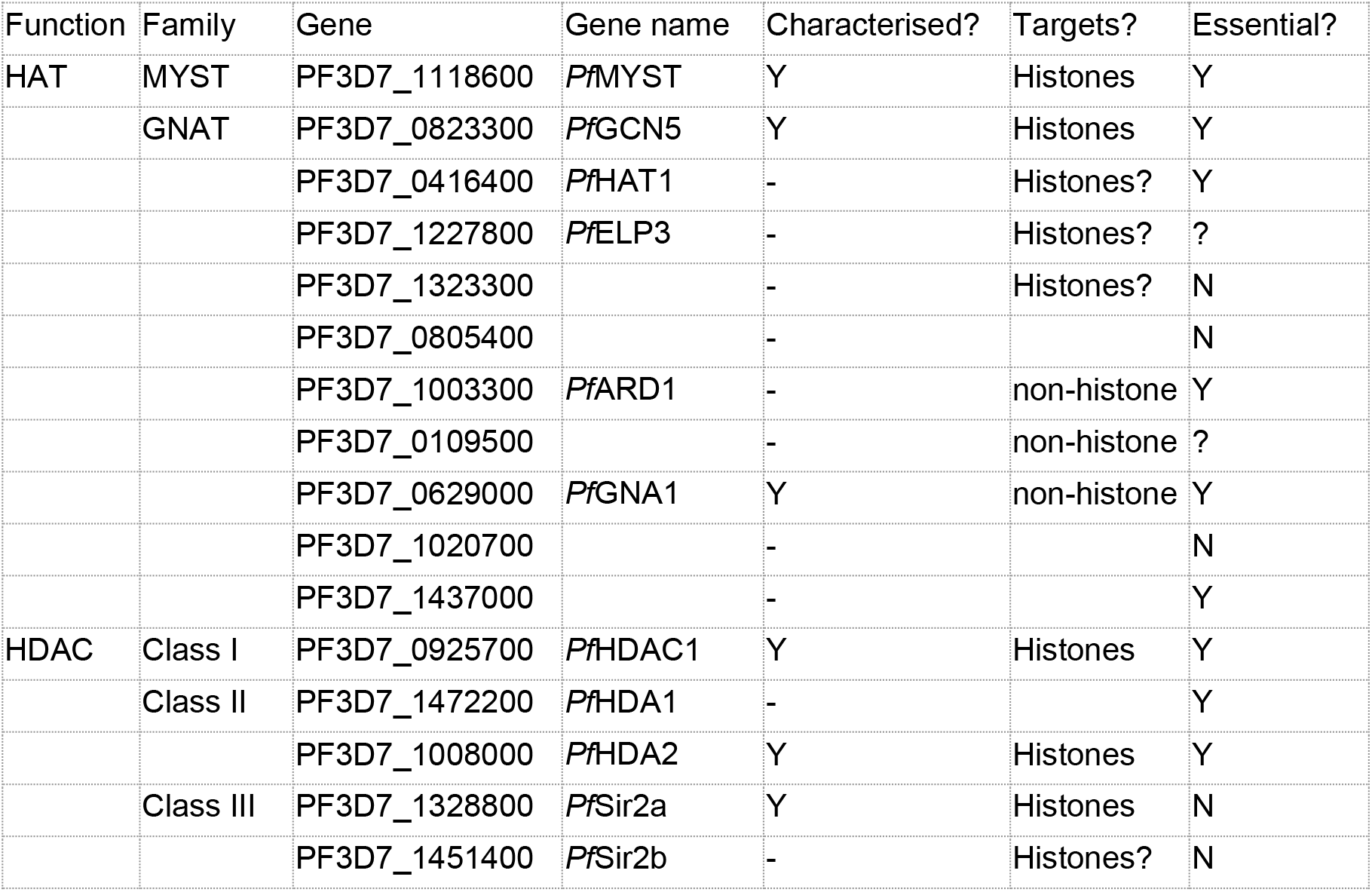
HAT and HDAC enzymes identified in *P. falciparum* Data assembled from published sources including [16, 17] and *plasmoDB*.*org*.

| Function | Family | Gene | Gene name | Characterised? | Targets? | Essential? |
| --- | --- | --- | --- | --- | --- | --- |
| HAT | MYST | PF3D7_1118600 | *Pf*MYST | Y | Histones | Y |
|  | GNAT | PF3D7_0823300 | *Pf*GCN5 | Y | Histones | Y |
|  |  | PF3D7_0416400 | *Pf*HAT1 | - | Histones? | Y |
|  |  | PF3D7_1227800 | *Pf*ELP3 | - | Histones? | ? |
|  |  | PF3D7_1323300 |  | - | Histones? | N |
|  |  | PF3D7_0805400 |  | - |  | N |
|  |  | PF3D7_1003300 | *Pf*ARD1 | - | non-histone | Y |
|  |  | PF3D7_0109500 |  | - | non-histone | ? |
|  |  | PF3D7_0629000 | *Pf*GNA1 | Y | non-histone | Y |
|  |  | PF3D7_1020700 |  | - |  | N |
|  |  | PF3D7_1437000 |  | - |  | Y |
| HDAC | Class I | PF3D7_0925700 | *Pf*HDAC1 | Y | Histones | Y |
|  | Class II | PF3D7_1472200 | *Pf*HDA1 | - |  | Y |
|  |  | PF3D7_1008000 | *Pf*HDA2 | Y | Histones | Y |
|  | Class III | PF3D7_1328800 | *Pf*Sir2a | Y | Histones | N |
|  |  | PF3D7_1451400 | *Pf*Sir2b | - | Histones? | N |

### Could histone lactylation be important more broadly in apicomplexan parasites?

The above discussion focusses on the principal human malaria parasite, *P. falciparum*. However, other *Plasmodium* species that are able to achieve high parasitaemias do cause hyperlactataemia, including the zoonotic macaque parasite *P. knowlesi* and the rodent model species *P. berghei* and *P. yoelii*. These were recently reported to raise blood lactate in mice to ∼12mM and 18mM respectively [18] – mimicking levels in severe human malaria. As in *P. falciparum* infections, this correlates with respiratory distress in human patients and with equivalent phenotypes in mice. Could all *Plasmodium* species therefore have evolved to sense blood lactate and modulate virulence phenotypes accordingly – and if so, which phenotypes would be modulated? The rodent species do not show classical cytoadherence and do not have *var* genes, but *P. knowlesi* has a gene family called *sicavar*, analogous to but technically different from the *var* gene family [19]. If *sicavar* genes were also affected by histone lactylation, this would imply an evolutionarily conserved pathway. All *Plasmodium* species could also share the epigenetic control of gametogenesis and stress-resistance.

Beyond the *Plasmodium* genus, related apicomplexan parasites (*Toxoplasma, Babesia, Cryptosporidium*, etc.) also have public-health importance. Only a few of these are blood-dwelling. They may reside in niches where lactate levels vary, and they may use epigenetics to control their biology, but none of them cause hyperlactataemia like *Plasmodium*. An initial protein ‘lactylome’ was recently published for *T. gondii*, identifying a wide variety of lactylated proteins including histones [20]. Whether or how these are functional, however, remains to be elucidated – in *Toxoplasma*, for example, the key stress-induced phenotypic switch from tachyzoite to bradyzoite is controlled by a transcription factor rather than an epigenetic switch [21]. Thus far, it appears that the biology proposed in this article might have evolved uniquely in *Plasmodium* due to the unique features of mammalian malaria.

## CONCLUSION

Histone lactylation is an entirely new epigenetic pathway in the protozoan parasite *Plasmodium*. It suggests a novel mechanism of host-parasite interaction in malaria: a disease in which the host frequently develops the potentially fatal complications of hyperlactataemia and respiratory distress. If malaria parasites have indeed evolved to sense and respond to this epigenetically then the implications for virulence and malarial disease are compelling.

#### OPEN QUESTIONS

- What is the full catalogue of histone lactyl modifications in *Plasmodium*? Are parasite-specific histone variants, particularly those associated with active genes, preferentially or uniquely lactylated? Are modified histone sites conserved across *Plasmodium* species?
- What are the temporal dynamics of histone lactylation after exposure to exogenous or endogenously-generated lactate?
- Which enzymes control histone lactylation in *Plasmodium?* Are ‘moonlighting’ HAT and HDAC enzymes entirely responsible?
- Where does histone lactylation occur throughout *Plasmodium* genomes? By combining chromatin immunoprecipitation (ChIP-seq) and RNA sequencing (RNA-seq), the gene types most affected by histone lactylation could be identified along with resultant changes in expression, thus pinpointing the likely biological roles for the epigenetic mark. Would the affected genes be similar in different *Plasmodium* species?
- Does histone lactylation control virulence phenotypes, such as cytoadherence, gametocytogenesis and stress-resistance – and if so, which target genes are responsible?
- In human malaria patients, is there a correlation between in-host hyperlactataemia, parasite virulence phenotype(s) and expression of lactyl-modified genes?

## ACKNOWLEDGEMENTS

I am grateful to Anders Jensen and Linda Onyeka Anagu for early experiments on this topic, and to Cameron Smith and Mike Deery for sharing and re-analysing unpublished mass spectrometry datasets.

The project (conceived in late 2019) has thus far received no specific funding; it will be funded from November 2022 by Wellcome Discovery Award 225171/Z/22/Z.

